# Innate floral object identification in the generalist solitary hoverfly pollinator *Eristalinus aeneus*

**DOI:** 10.1101/2024.02.08.579518

**Authors:** Aditi Mishra, Anupreksha Jain, Padmapriya S Iyer, Ashwin Suryanarayanan, Karin Nordström, Shannon B. Olsson

## Abstract

All animals must locate and identify food for their survival. Most insects are solitary. Newly emerged solitary insects must therefore employ innate identification of food cues to locate relevant nutritive objects from a distance. Innate preferences for food cues should be both specific enough to allow discrimination between food and non-food objects and general enough to allow for the variety of food objects relevant to each species’ ecology. How insects with small nervous systems are able to encode these cues innately is an area of intense study for ecologists and neuroscientists alike. Here, we used the solitary generalist pollinator *Eristalinus aeneus* to understand how the innate search template used to identify multiple floral species can arise through a small number of sensory cues spanning multiple modalities. We found that innate floral choices of the hoverfly *E. aeneus* are a product of contextual integration of broad, plant-based olfactory cues and visual cues, where a combination of radial symmetry and reflection in the 500-700 nm wavelength range was particularly important. Our study, therefore, shows how tiny brains can efficiently encode multimodal cues to identify multiple relevant objects without prior experience.

## Background

Identifying objects is fundamental to animal life. All organisms must, for instance, be able to distinguish objects of nutritive value from non-nutritive objects. Identifying food in the high-dimensional natural world is particularly challenging. For many organisms, objects can be learned individually or socially over their lifetime. However, in some cases, the innate identification of certain objects is crucial for survival. Several vertebrates innately identify cues associated with predators (Hawkins et al. 2004), kin (Rosa-Salva et al. 2018), and invertebrates like arachnids are innately attracted to their prey (Dolev and Nelson 2014).

Most insects are solitary, and therefore the majority of insect species must identify initial food objects without any social learning or larval experience with food as found in social insects (Farina et al. 2005; Arenas et al. 2009). Essentially, solitary insects can employ three sensory strategies to discriminate between food and non-food objects innately. First, the insect can accept a large set of objects as food with little discrimination. Second, the insect can employ a minimal set of cues common to all nutritive objects. Third, the insect can use a specific combination of sensory cues signifying a particular object or set of objects.

Attraction to many objects with little discrimination is computationally simple, but metabolically and energetically challenging, as it requires exploring several objects and a potentially low-quality diet. The increased search and handling cost, coupled with the low rewards, would render this strategy unviable for most organisms. Being attracted to a single or small and specific set of sensory cues, such as rare volatiles, has been observed in many specialist insects such as bees (Milet-Pinheiro et al. 2013), flies (Cha et al. 2011), aphids (Chapman et al. 1981), and weevils (Bartlet et al. 1997). However, a narrow preference could be inefficient when those particular resources are scarce, necessitating a generalist diet that can utilize a variety of resources as food.

Generalist solitary insects thus face a two-fold problem. First, they must discriminate nutritive objects from the clutter of the natural world. Second, these insects should ideally generalize food to multiple objects. The innate preferences of generalist solitary insects must therefore be flexible to accommodate varied resources yet maintain high accuracy. Thus, as a first principle, one would hypothesize that innate food identification in solitary generalist insects should be guided via attraction to a small set of cues indicative of several relevant food objects. In this study, we test this hypothesis using a solitary insect generalist pollinator, the hoverfly *Eristalinus aeneus* (Scopoli, 1763), and examining the minimal set of floral cues necessary for them to innately identify floral objects.

Long-distance floral choice in generalist insect pollinators is mediated by several factors including olfactory cues (Andersson and Dobson 2003; Primante and Dotteral 2010), and visual cues such as hue which is the peak of reflectance spectra (An et al. 2018; Trunschke et al. 2021), saturation signifying the spread of the reflectance spectra (Lunau 1990; Trunschke et al. 2021), brightness, which is intensity or amplitude of reflectance spectra (Free 1970; Trunschke et al. 2021), and shape (Free 1970; Howard et al. 2019). Pollinators are known to utilize combinatorial signatures for foraging (Riffell and Alarcon 2013; Howard et al. 2019; Balamurali et al. 2020; Matoušková et al. 2023) instead of single cues to identify floral objects. Further, cross-modal interactions can alter the valence of a stimulus. For example, colour modulates proboscis extension reflexes (PER) to learnt odours in bees (Mota et al. 2011), and ovipositional odours decrease preference to specific colours in female pierid butterflies (Balamurali et al. 2020). These findings indicate the need to study the floral object as a whole. Hence, multimodality and multidimensionality of experimental stimuli is key to understand the innate floral search template from both ecological and physiological perspectives. However, while we know the innate preferences of some pollinators to specific floral cues, pollinator behaviour in response to combinations of complex floral cues has been understudied in solitary insects. Most studies have occurred in social hymenopterans where innate attraction is difficult to discern because of social learning from the hive (Farina et al. 2005; Arenas et al. 2009).

Hoverflies are important solitary non-bee pollinators that visit at least 72% of global food crops (Doyle et al. 2020). They exhibit colour vision (Ilse 1949), with a preference for yellow hue in human colour vision in some hoverfly species like *Episyrphus balteatus* (Sutherland et al. 1999) and *Eristalis tenax* (An et al. 2018). However, hoverflies also visit flowers with other colours, and use multimodal cues for foraging in the wild (Lunau 2014; Nordström et al. 2017). Since the sensory physiology (Lunau 1990, Lunau 2014, An et al. 2018), ecological niches (Rotheray and Gilbert 2015; Nordström et al. 2017), and attractive floral signatures (Nordström et al. 2017) are known for hoverflies, they make excellent model organisms to investigate the innate cues used to identify objects in generalist insects. Further, the solitary nature of hoverflies (Rotheray and Gilbert 2015) removes confounding effects of social learning from the hive (Farina et al. 2005), and sensitivity (Arena et al. 2009) or floral biases (Farina et al. 2005) arising therein.

In this study, we assessed the minimal set of floral cues used by flower-naïve *E. aeneus* to identify floral objects. As stimuli, we employed a multimodal floral lure derived from several flowering species attractive to hoverflies across continents in the field (Nordström et al. 2017)). By artificially replicating these multimodal cues composed of floral colour, shape, size, and odour, the previous study found that hoverflies of multiple species and in multiple environments were attracted to a small set of cues common to several flower species, implying that these cues could be innately attractive (Nordström et al. 2017). We tested the innate attractiveness of this small set of cues to flower-naïve *E. aeneus* by presenting them with sequential perturbation of these cues. Evaluating the innate preferences of hoverflies helps us understand how this important pollinator identifies flowers in the natural world, with implications for our understanding of complex object identification in small insect brains.

## Materials and methods

### Study design

The floral characteristics tested in the current study were first derived from a cross-continental field study using wild hoverflies from the environment (Nordström et al. 2017). This previous study found that models with a specific combination of visual and olfactory cues (henceforth referred to as the positive control, or v+o+) were ubiquitously attractive to hoverflies across multiple ecosystems without any nutritive rewards (Nordström et al. 2017). This preference across environments implies that v+o+ could be innately attractive to flower naïve hoverflies. To test this hypothesis, the current study conducted two choice assays with freely flying, flower-naïve *E. aeneus* to examine (1) if the v+o+ model was innately attractive as well, and, if so, (2) which visual or odour characteristics are important for innate attraction. In the previous field study, the negative control was a dark circular disk with no odour, and it received significantly fewer visits (Nordström et al. 2017). Thus, in our study we chose a similar negative control (v-o-) of equivalent diameter to the positive control, without any odour. To control for confounding environmental factors, all experiments were performed in lab conditions in a black cage with a black background, black mesh sides, and checker board top (Supplementary Fig. S1). Preliminary tests indicated that an arena with this configuration increased the response rates of tested hoverflies compared to green or white arenas of the same size.

In the field study, wild, free-ranging hoverflies were observed to land sparingly, with the majority of responses being directed flights to within 15 cm of the model, after which the hoverflies flew away (Nordström et al. 2017). Hence, in our study we similarly defined a response as either landing from above or directed flights within 5 cm towards the model in our arena (see “Two-choice assays”). To determine which visual and odour cues were important for attraction, we adopted a reductive approach where, unless stated otherwise, *E. aeneus* were provided a choice between v+o+ and a model with perturbation to one feature of the v+o+, with all other features kept constant.

## Artificial floral models

### Visual cues of the artificial floral models

The artificial floral models were printed using WoL 3D printer silk white PLA filament (WOL3D, India) and Raised N2 3D printer (Raise3D, USA), ensuring that they were identical. The visual and olfactory cues of the positive control (v+o+) had been previously standardized (Nordström et al. 2017) as a radially symmetric, petaloid, non-nutritive object measuring 36 ± 0.1 mm in diameter with the blend of five floral volatiles and high reflectance in medium wavelengths (500 – 700 nm). This model appears yellow hued in human colour vision.

The floral models were painted with Camlin acrylic paints (KOKUYO CO., LTD.; Japan). A white base coat was provided to all floral models, followed by specific ratios of paints mixed to achieve the needed reflectance spectrum (Table S1). All visual models, and their reflectance spectrums are shown in Fig. 1 (see Table S1 for full details). The surface area of all models except negative control is the same, the longest diagonal of Hexagon+ (∼32 ± 0.1 mm) is smaller than the diameter of v+, while the major chord of Asymmetric+ (∼40 ± 0.1 mm) is longer than diameter of v+. The four petals of asymmetric model are one identical ellipse of minor chord of 6.8 mm, and major chord 40 mm rotated at 0°, 45°, 200°, 285° making it asymmetric. Hoverflies, known to detect differences smaller than 1° while moving (Nordström and O’Carroll 2006), should be able to perceive this asymmetry.

**Fig. 1.**
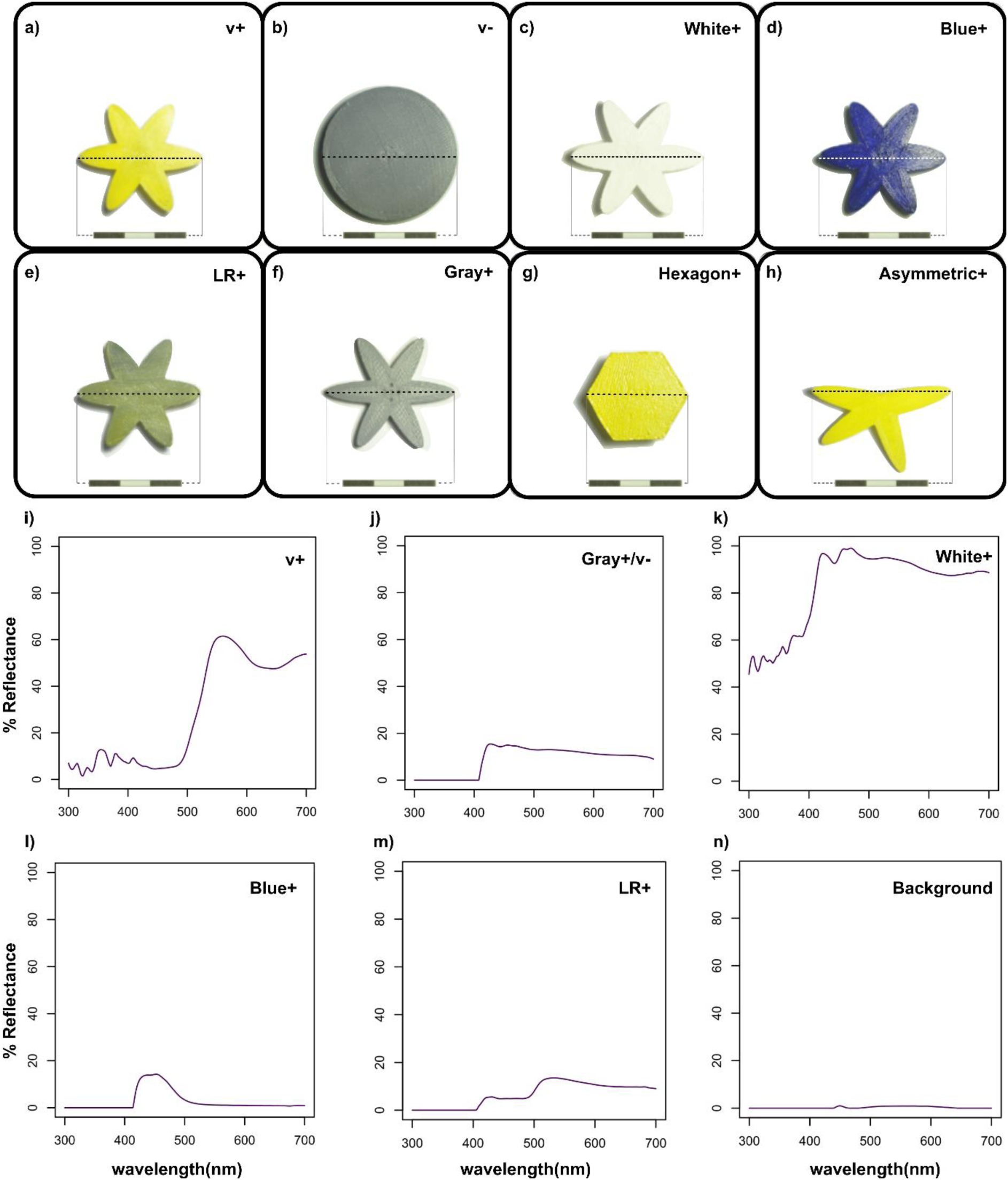
Different visual models used in the study and their respective reflectance spectra. a), i) positive visual control (v+), b), j) negative visual control (v-), c), k) broad-spectrum model (White+), d), i) Blue+, e), m) low reflectance model (LR+), f), i) broad-spectrum low reflectance model (Gray+), g), a) undisrupted model (Hexagon+), h), a) asymmetric model (Asymmetric+). n) background of the experimental arena. Scale bar 3 cm.

The reflectance of all models was measured using a portable spectrophotometer (Ocean optics Jaz-200 spectrophotometer with the optic fiber tip of 25-cm, 600-μm Premium Fiber, UV/Visible, FL, USA), after calibrating the spectrophotometer to a white balance (Labsphere certified reflectance standard USRS-99-010 AS-01158-060 S/N 99AA08-0615-3676) under the 5000 Lux LED lights used in the experimental setup (ORFLD-30W, Chromatic temperature 6500 K close to CIE standard illuminant D65 with a chromatic temperature of 6504 K). Output was the spectral power distribution curve of light reflected off an object (in percent reflectance) normalised to the white balance.

### Odour Cues of the artificial floral models

Models with o+ in their name have the same odour blend of five floral volatiles derived from cues attractive in field conditions across multiple continents (2-Ethyltoluene, p-Cymene, Undecanal, r-Limonene, 6-Methyl-5-hepten-2-one [Fluka], in mineral oil; Nordström et al. 2017). Blends were provided by pipetting 2 µl of the blend on grade 2 Whatman filter paper (1 cm x 5 cm) affixed to the artificial floral models. Refer to Table S2 for full details.

Emission rates for o+ were calibrated empirically against measured release rates of component volatiles from real flowers in the field (Nordström et al. 2017; Table S3). The volatile emission from the models was assessed using a 100 µm PDMS solid-phase microextraction (SPME) fiber, model no:548575-U, SUPELCO, USA. The fiber was placed within 10 cm of the models with volatile blends in a foil covered, cleaned and heated 500 ml beaker for 30 seconds. The fibre was injected into the inlet of an Agilent 7890B gas chromatograph in splitless mode, to desorb volatiles at 270°C. The volatiles were then separated using an HP-5 MS column (30 m × 0.25 mm i.e., 0.25 µm film thickness) coupled with a 5977A MSD mass spectrometer. The column oven was kept at 40°C for 1 min, increased to 150°C at 25°C/min and then to 270°C at 100°C/min and held for 5 minutes. Compounds were identified by mass spectroscopy using Masshunter Qualitative Analysis software (B.07.00) and the mass spectra library NIST (National Institute of Standards and Technology and libraries created from authentic standards). The models were sampled at 5-minute intervals to check for contamination and determine release rates as a function of time. This enabled us to ensure similar release rates, retention times and absence of any contaminants during the behavioural assays (Fig. S2, Table S3).

### Two choice assays by freely-flying insects

Insects were reared as stated in Supplementary material S1. All tests were performed in a black Nylon mesh cage of side 50 cm (width x height x length: 50 cm x 50 cm x 50 cm) with a solid black base as background for the models, and a checker board roof under two 100 W flicker-free white LED flood lights. The illumination of the experimental arena was 5000 ± 50 Lux (mean ± s.e.) as measured using a lux meter (LX-101A HTC instruments). Two models were provided in the centre of the arena – 20 cm apart from each other, in the middle of the cage, 25 cm away from the front and the back of the cage. The model’s stalks were 20 cm high. Directly on top of the floral models, an exhaust duct was installed to ensure minimal contamination of the odour from the other model. The flies were captured individually using 50 ml Falcon tubes (Tarsons, catalog number: 500041), in which they were carried to the experimental arena (1 fly per tube), within 5 minutes. Each fly was placed on the floor of the experimental arena at the mid-point of the line joining the base of the stalks of the two models, on the cap of the falcon tube in which it was transported to the arena (Fig. S1). The set-up and a behavioural assay are shown in Online resource 2 and Fig. S1.

All observations were conducted in real time, with the observers always sitting across from the transparent front panel of the experimental arena. The observer’s line of sight was at the same plane as the models. A response was defined as landings or approaching at least 5 cm above the models, as per other studies (Nordström et al. 2017; Matoušková et al. 2023; Chapman et al. 2023). The time period for each observation was capped at 5 minutes. Only the first response by a hoverfly was counted, with the total number of hoverflies responding as described above as n_responder_. The testing arena was controlled for side bias by periodically switching the floral models. In the side bias assay, the positive control (v+o+) was placed at both sides.

### Electroantennography (EAG)

*E. aeneus* between the ages of 4 – 10 days were used for EAG and prepared as previously described (Batra et al. 2019). Data was acquired using EAG2000 software and insects were exposed to air, solvent blank, and volatile concentrations ranging from 10^-1^ g/ml to 10^-8^ g/ml presented in 10 μl aliquots on filter paper inserted into Pasteur pipettes as in (37). Data were acquired using Syntech intelligent data acquisition controller (IDAC-2, SYNTECH, Netherlands) and the stimuli was presented as pulses of 0.5 s for 10 seconds of recording using a custom-built odour delivery system, standardized as previously described (Batra et al. 2019). The electrophysiological response of flies was calculated as voltage deflections in millivolts (mV) as follows:

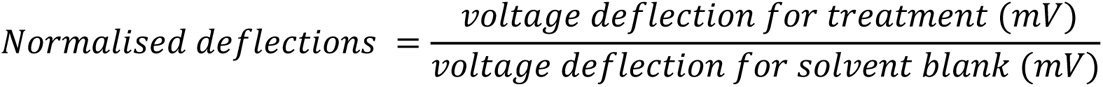

## Statistical analysis

For each assay, a series of Binomial GLMMs (Generalized Linear Mixed Models) were performed to test for differences in responses to the two models. These GLMMs were used to test: (1) whether flies responded differently to the two models in the different assays, (2) whether the proportion of responders out of the total number of flies tested in an assay, depended on the sex of the fly, and (3) whether sex had an effect on the preference of one model over the other for n_responder_ flies. More details of the GLMMs used are given in Supplementary Material 1. GLMMs were performed using the *glmmTMB* package (Brooks et al. 2017) in R version-4.0.5 (R team R.D.C.T 2014). Model diagnostics were checked using the *DHARMa* package (Hartig 2020), and model-predicted means and confidence intervals were extracted using the *emmeans* package (Lenth 2022), for additional information refer to Supplementary material S2.

Phi coefficient (ϵ) was used to estimate the effect size of the difference, calculated as:

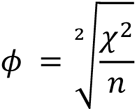

ᵡ^2^is the chi square value. Phi-coefficient is a special case of Cramer’s V, used to measure the effect size of associations among categorical variables, where a value of 0 signifies no effect, 0.1 a very small effect, and 0.5 a large effect (Thimmegowda et al. 2020). Preference index for a model in an assay was defined as the percentage of response received by a model in a two-choice assay, calculated as:

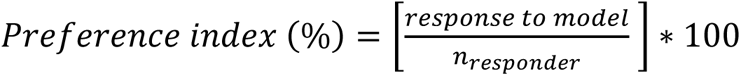

## Results

### Assessing the innate attractiveness of artificial floral objects

We tested side bias in the arena by placing v+o+ on both sides and found no significant side bias to either side of the arena (n_responder_ = 105; Preference index to v+o+ on left = 48.54, p = 0.768 Fig. 2a.1). To investigate whether cues attractive to hoverflies in the field (Nordström et al. 2017) were innately attractive to naïve *E. aeneus*, hoverflies were given a choice between positive control (v+o+, Fig. 1a) and negative control models (v-o-, Fig. 1b) with similar cues as tested in the field (Nordström et al. 2017). The response to v+o+ the positive control was significantly higher than to the negative control (n_responder_ = 96, preference index to v+o+ = 73.95, p = 7.18**\***10^-6^, phi = 0.35; Fig. 2a.2). This suggests that flower naïve *E. aeneus* are innately attracted to the positive control (v+o+), as suggested by previous field experiments (Nordström et al. 2017).

**Fig. 2.**
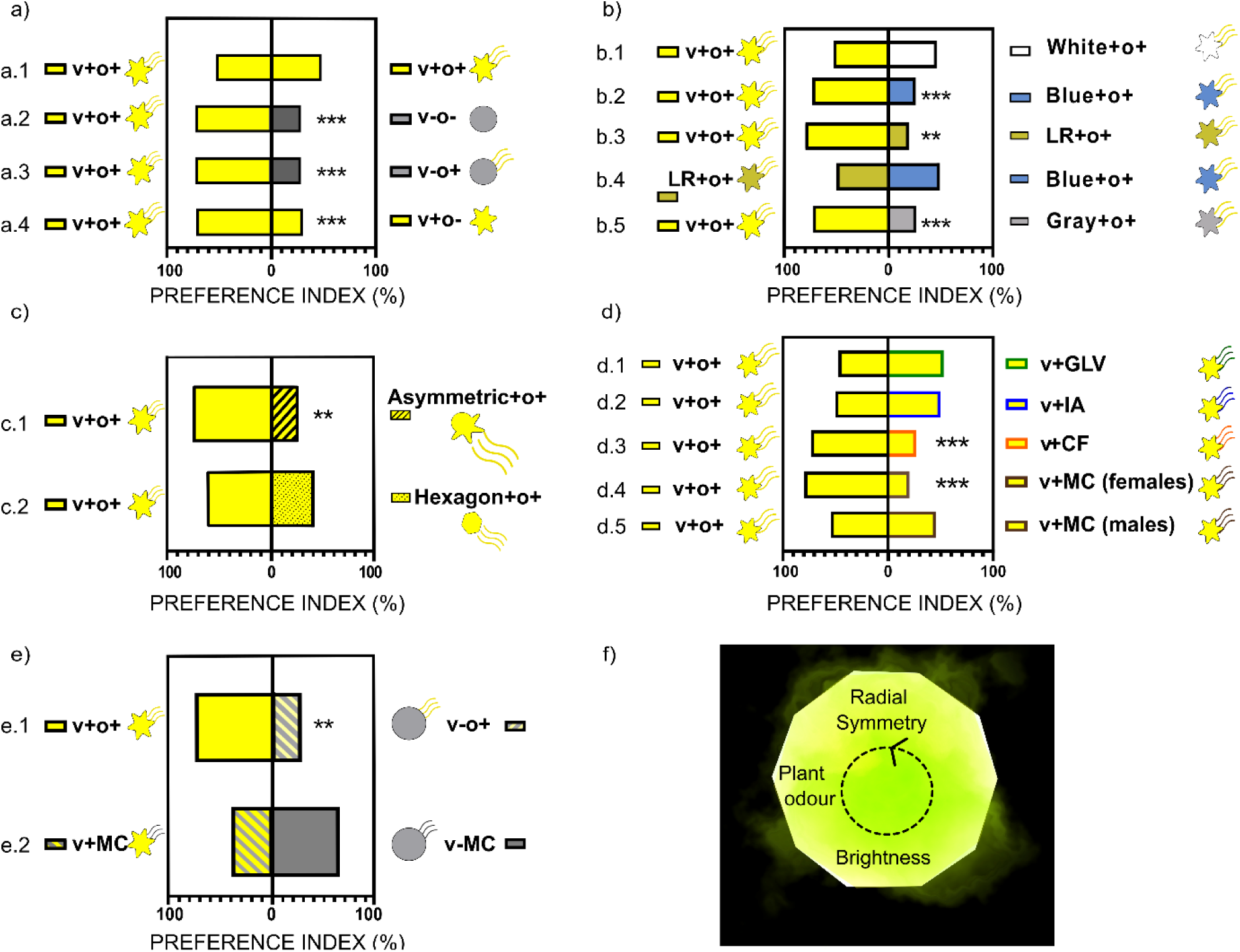
Preference indices of naïve *E. aeneus* when given a choice between the positive control (v+o+) against various single cue perturbations as listed. Stars indicate significant differences based on Binomial GLMM (* for p < 0.05, ** for p < 0.01, *** for p < 0.001; Table S4). (a) Assays for multimodality of positive control v+o+ against (a.1) itself (side bias assay), (a.2) negative control, v-o-, (a.3) negative visual control with floral odour, v-o+, (a.4) positive visual control with no floral odour, v+o-; (b) Assays for hue and reflectance of positive control v+o+ against (b.1) broad-spectrum model White+o+, (b.2) Blue+o+, (b.3) low reflectance model LR+o+. (b.4) low reflectance model LR+o+ against Blue+o+, and (b.5) positive control v+o+ against low reflectance broad spectrum model Gray+o+; (c) Assays for shape of positive control v+o+ against (c.1) Asymmetric+o+ and (c.2) undisrupted Hexagon+o+; (d) Assays for odour of positive control v+o+ against (d.1) v+ with green leaf volatile v+GLV, (d.2) v+ with isoamyl acetate v+IA, (d.3) v+ with coffee furanone v+CF, (d.4) v+ with meta-cresol v+MC (female flies only) (d.5) v+ with meta-cresol v+MC (male flies only). (e) Female flies given a choice between (e.1) positive control v+o+ and negative visual control with odour cues v-o+, and (e.2) positive visual control with meta-cresol (v+MC) and negative visual control with meta-cresol (v-MC). Pictograms indicate visual cue presented and wavy lines indicate odour with different colours for different odour identities. (f) Theoretical model of minimal set of floral cues composed of radial symmetry, brightness and plant odours necessary to attract response from a naïve hoverfly

### Two choice assays between multimodal and unimodal floral objects

Some studies report hoverflies like *Episyrphus balteatus* can use unimodal cues for identifying flowers (Primante and Dotteral 2010). To evaluate if either odour or visual cues alone were sufficient for *E. aeneus* to be attracted to our floral model, naïve *E. aeneus* were given a choice between the positive control (v+o+) and a negative visual model with floral odour (gray disk; v-o+) or the positive visual model without floral odour (v+o-). We found that the hoverflies responded to the positive control in significantly greater numbers over either unimodal model (v-o+: n_responder_ = 90, preference index to v+o+ = 73.33, p = 2.2**\***10^-5^, phi = 0.357; Fig. 2a.3; v+o-, n_responder_ = 108, preference index to v+o+ = 70.37, p = 4.05**\***10^-5^, phi = 0.296; Fig. 2a.4). This indicates that both odour and visual cues are important for the innate attractiveness of the positive control v+o+.

## Parsing visual cues

### Two choice assays between floral objects of different colours

The colour of any object is impacted by its hue (peak of reflectance spectra), saturation (spread of reflectance spectra), and brightness (amplitude of reflectance spectra). Several generalist pollinators show preferences for specific colour hues in lab settings (Guirfa et. al 1995; An et al. 2018) and preference for saturated colours has been reported in bees (Lunau 1990). In all assays parsing visual cues, all the visual morphs were paired with the positive control odour o+ as indicated in their labels.

*E. aeneus* were given a choice between the positive control (v+o+) and a Broad-spectrum model (White+o+, see Fig. 1c). The positive control v+o+ reflects across wavelengths from 500 nm to 700 nm, with a peak around 565 nm (Fig. 1i). The White+ model reflects light across a broader range of wavelengths (Fig. 1k). Hoverflies responded to both models equally (n_responder_ = 55; preference index to v+o+ = 52.73, p = 0.686, Fig. 2b.1). Thus, the response in flower naïve hoverflies to our flower models is not restricted to objects with only yellow hues. *E. aeneus* were then given a choice between the positive control and Blue+o+ (Fig. 1d, Online resource 2). The Blue+o+ model has a peak at 440 nm, and reflectance in the 400 – 500 nm range (Fig. 1l). However, the Blue+ model reflected very little around the wavelengths found in the positive control (500 – 700 nm; Fig. 1i). We found that hoverflies respond to v+o+ significantly more than the Blue+o+ model (n_responder_ = 64, preference index to v+o+ = 73.44, p = 0.000327, phi = 0.33; Fig. 2b.2). This suggests that floral objects with yellow hues are preferred by flower-naïve hoverflies over objects with blue hues. This is similar to observations in other flower-naïve hoverflies like *Eristalis tenax* (Matoušková et al. 2023).

### Two choice assays between floral objects with different brightness

Generalist pollinators like bumblebees have been reported to use both hue and achromatic contrast (brightness) for flower choice (Giurfa et al. 1995). Since the background and the illumination was constant in our assay, we lowered the amplitude of reflectance of our models to study the impact of brightness. LR+o+ (Fig. 1e) has lower brightness than v+o+ but the same reflectance peak or hue as the positive control v+o+ (Fig. 1m, 1i). We found that *E. aeneus* responded to the positive control significantly more than to the LR+o+ model (n_responder_ = 30, preference index for v+o+ =80, p = 0.00239, phi = 0.5; Fig. 2b.3, Online resource 2). This suggests that floral objects with higher contrast or brightness are preferred by flower-naïve hoverflies.

We then presented *E. aeneus* with LR+o+ against Blue+o+. In contrast to the above assay, the flies in this assay responded to both floral models in equal proportions (n_responder_ = 64; preference index for LR+o+ = 50, preference index for Blue+o+ = 50, p = 1, Fig. 2b.4). models. This lack of preference indicates that both brightness and hue play a role in the response of flower-naïve hoverflies to floral objects.

*E. aeneus* were also given a choice between the positive control (v+o+) and a low reflectance broad spectrum model Gray+o+ (Fig. 1f, 1j). The hoverflies responded to the positive control in significantly higher numbers (n_responder_ = 56, preference index for v+o+ = 73.21, p = 0.000862, phi = 0.3; Fig. 2b.5). This is in contrast to the equal proportion of responses observed between the positive control and the higher reflectance broad spectrum model (White+o+; Fig. 2b.1), further indicating the importance of brightness in response of flower-naïve hoverflies.

### Two choice assays between floral objects of different shapes and symmetry

Many insect pollinators like bees prefer symmetry and disruptive shape as a floral cue (Free 1970), and some hoverflies like *Eristalis tenax* prefer both radially or bilaterally symmetric objects equally (Matoušková et al. 2023). Hence, we next evaluated the preference for symmetry and disruptive shape in our floral models. *E. aeneus* exhibited a preference for the positive control (v+o+) against an asymmetric model (Asymmetric+o+, Fig. 1h; Fig. 2c.1; n_responder_ = 61, preference index to v+o+ = 75.41, p = 0.00164, phi = 0.382). However, *E. aeneus* responded to the v+o+ and undisrupted model equally (Hexagon+o+, Fig. 1g; Fig. 2c.2; n_responder_ = 65, preference to v+o+ = 60, p = 0.181). All shape perturbation models had the same surface area as the positive control and all our models were presented horizontally. This suggests that symmetry, but not necessarily disruptive shape, is important for innate floral object preference in *E. aeneus*.

## Parsing odour cues

### Two Choice assays between floral objects of different odour identities

Odour alone can be sufficient for object identification in some hoverflies (Primante and Dotteral, 2010). To evaluate the relevance of floral odour identity, *E. aeneus* were given a choice between the positive control (v+o+) and a positive visual model with the vegetative odour cis-3-hexenyl acetate. Cis-3-hexenyl acetate, also known as a green leaf volatile (v+GLV), is present in many floral scents (Effmert et al. 2005; Cozzolino et al. 2015; Nordström et al. 2017; Feng et al. 2017) and has been reported to increase pollinator attraction (Cozzolino et al. 2015). While parsing odour cues, we kept the associated visual cues across the morphs same as the positive control visual cues (v+). We found that flies responded to both models equally (n_responder_ = 61; preference index for v+o+ = 47.54, p = 0.701, Fig. 2d.1). Further, *E. aeneus* responded equally to the positive control and a positive visual model with the fruit odour isoamyl acetate (n_responder_ = 59; preference index for v+o+ = 48.28, p = 0.696 Fig. 2d.2).

Given that *E. aeneus* did not exhibit any preference between models with either floral, vegetative, or fruit odour, we next evaluated if the presence of any odour was sufficient for attraction. The plant-derived compound 2-methyltetrahydrofuran-3-one, also known as coffee furanone, is known to repel many insects including dipterans like *Drosophila melanogaster* and *Aedes aegypti* at a 10^-1^ g/ml concentration (Batra et al. 2019). In our assay, we found that *E. aeneus* responded to the positive control in significantly higher numbers than a positive visual model with coffee furanone (v+CF, n_responder_ = 60, preference index to v+o+ = 73.33, p = 0.00053, phi = 0.33; Fig. 2d.3). In summary, *E. aeneus* is attracted to a wide range of plant odours when coupled with visual cues in floral objects, but the presence of any odour is insufficient for attraction. Note that all assays were conducted using concentrations similar to empirically measured release rates of odours from flowers in nature (Nordström et al. 2017) (Fig. S2, Table S3, see Methods for details).

### Contextual pairing of odour and visual cues

We examined if *E. aeneus* innately prefer relevant odours irrespective of context. Herbivore dung piles are an important oviposition site for hoverflies (Rotheray and Gilbert 2015), and meta-cresol is an odour abundantly found in herbivore dung (Nair et al. 2018). Thus, unlike the other plant-derived odours tested, meta-cresol is more likely to be an odour with neutral or even positive valence in a non-food (oviposition) context. We examined the preference of *E. aeneus* between the positive control and a positive visual model with meta-cresol (v+MC). Interestingly, we observed sexually-dimorphic response rates (ratio of n_responder_ to total number of flies tested) and responses in this assay, with a statistically significant impact of sex on both response rates (Table S5) and responses (Table S6).While female *E. aeneus* hoverflies responded significantly more to the positive control (v+MC, n_responder_ = 115, preference index for v+o+ = 80.87, p = 1.2**\***10^-9^, phi = 0.42; Fig. 2d.4), male *E. aeneus* responded equally to both models (n_responder_ =126; preference index for v+o+ = 51.59, p = 0.469 Fig. 2d.5).

To assess whether the female-specific response to meta-cresol was dependent on the visual context provided, we presented mated female hoverflies with a choice between a negative visual model paired with meta-cresol (v-MC) against the positive visual model and meta-cresol (v+MC). In contrast to behaviour of females to models presented with floral odour (Fig. 2e.1), mated females responded more to the negative visual control when paired with meta-cresol (v-MC) vs. the positive control (v+MC), but the difference was not statistically significant (n_responder_ = 50; preference index for v+MC = 40, preference index for v-MC = 60, p = 0.16, Fig. 2e.2).

Sexual dimorphism was not observed when v+ was paired with other the volatiles (o+, IA, GLV or CF; Table S5, Table S7). Note that the sexually dimorphic response to meta-cresol was not due to differences in chemoreception as there was no statistically significant difference in the sensitivity of male and female *E. aeneus* to meta-cresol since both males and females start detecting meta cresol at the concentration of 10^-2^ g/ml (n_male_ = 14, n_female_ = 18, Fig. S3, Table S8, Table S9). Fig. 2f shows the hypothesized floral model which should be sufficient to elicit innate floral object identification in hoverflies.

### Two choice assays between floral objects of different odour concentrations

Generalist pollinators have been documented to utilize the intensity of floral volatiles as a cue to identify flowers (Wright and Schiestl 2009). To understand if the intensity of floral odour was relevant for innate choices, *E. aeneus* were given a choice between the positive visual control (v+) and an odour blend that had been diluted to 1% of the original loading concentration calibrated to concentrations measured from real flowers (Nordström et al. 2017). No significant preference was observed between the 1% positive odour model (v+(1%o+)) and a model with GLV odour (v+GLV; n_responder_ = 61; preference index for v+(1%o+) = 40.98, preference index for v+GLV = 59.02, p = 0.161, Fig. 3a.1) the v+1%GLV vs. v+(1%o+) (n_responder_ = 30; preference index for v+(1%o+) = 43.33, preference index for v+(1%GLV) = 56.67, p = 0.467, Fig. 3a.2), or the positive control v+o+ vs. a 1% GLV model (v+(1% GLV); (n_responder_ = 57; preference index for v+o+ = 59.65, preference index for v+(1%GLV) = 39.35, p = 0.148, Fig. 3a.3). A significant difference in preference index was observed between models containing the innately attractive odour blend at 100% or 1% concentration (v+o+ against v+(1%o+); n_responder_ = 62, preference index for v+o+ = 64.52, p = 0.0243; Fig. 3a.4).

**Fig. 3.**
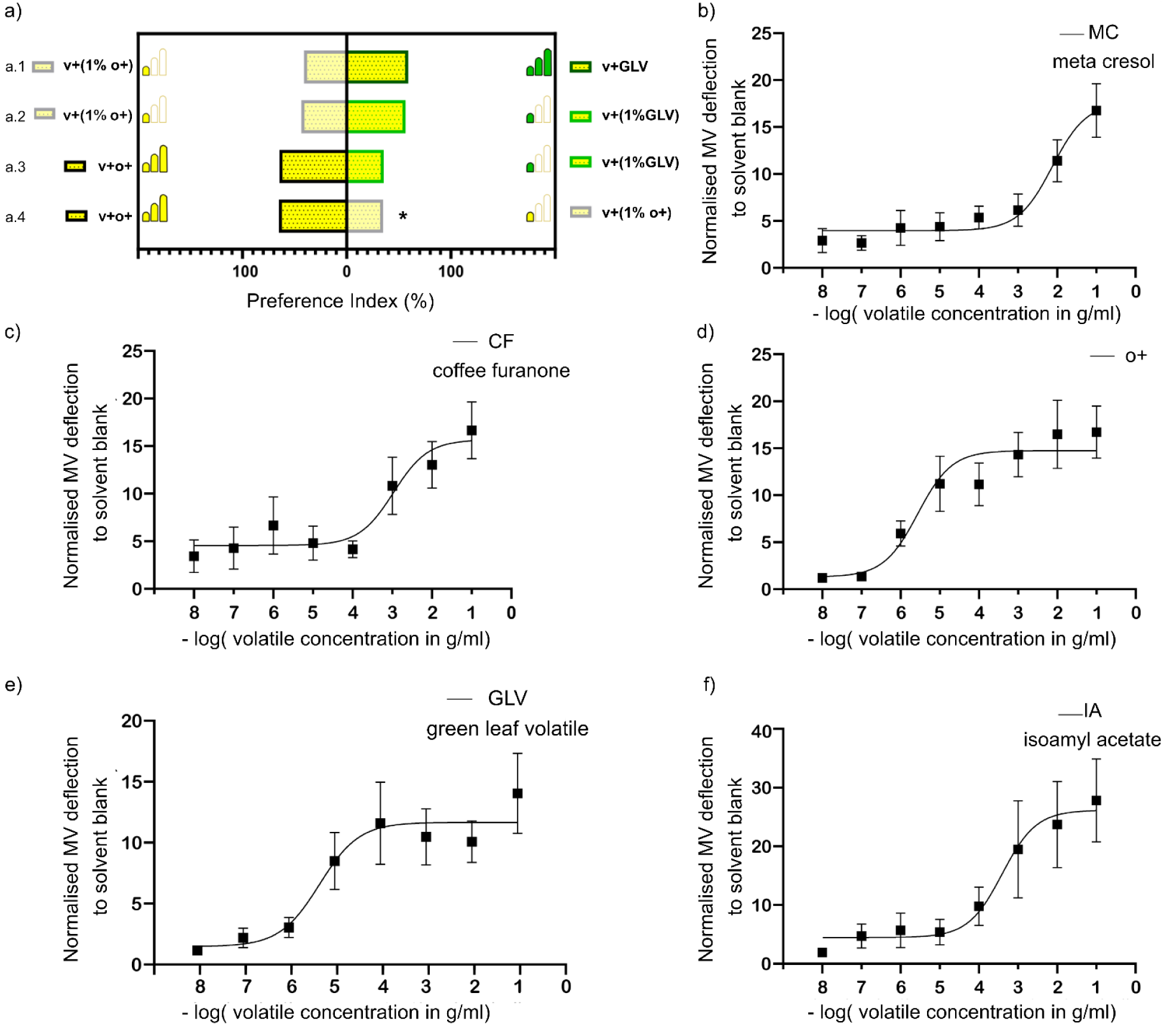
Preference indices of naïve *E. aeneus* when given a choice between floral objects with positive visual control v+ and different concentrations of odour cues, as well as electroantennogram responses to varying concentrations of these different odour cues ranging from 10^-8^ g/ml to 10^-1^ g/ml. Stars indicate significant differences based on Binomial GLMM (**\*** for p < 0.05, **\*\*** for p < 0.01, **\*\*\*** for p < 0.001; Table S4). (a) Preference index of naïve *E. aeneus* when given a choice between (a.1) the positive visual control v+ containing 1% positive odour blend (v+1%o+) against the positive visual control v+ containing GLV (v+GLV); (a.2) the positive visual control v+ containing 1% positive odour blend (v+1%o+) against the positive visual control v+ containing 1% GLV (v+ 1%GLV); (a.3) positive control (v+o+) against v+(1%GLV); (a.4) positive control (v+o+) against positive visual control with 1% positive odour blend (v+1%o+). Pictograms indicate the volatile concentration; one filled bar indicates diluted 1% volatiles. (b-f) Concentration-dependent EAG response curves (least-squares fit with mean and standard error bars, n≥20) shown for (b) meta-cresol (MC) n = 34, (c) coffee furanone (CF) n = 20, (d) positive odour blend (o+) n =22, (e) green leaf volatile (GLV) n =27, and (f) isoamyl acetate (IA) n = 20, all normalized against the solvent control – hexane

Since the naïve hoverflies used in this study did not show any behavioural preference towards different concentrations of odours of GLV and o+, except when o+ was presented at 1% of its original concentration, we additionally conducted electroantennograms (EAGs) to assess the antennal detection of odours at these concentrations. The EAGs indicated that the hoverflies detected all the volatiles at the loading concentrations as used in the behavioural studies (n ≥ 20, Fig. 3b-g). Further, we found that the concentration-response curves for the positive odour blend o+ (Fig. 3d), and for green leaf volatile GLV (Fig. 3e) plateaued at a 10^-5^ g/ml range.

## Discussion

Generalist insects face a complex dilemma in identifying nutritive sources. To locate viable resources, they must generalize a large pool of objects with diverse sensory cues to the class of nutritive objects, while rejecting irrelevant objects that could share some of these attributes. Our results indicate that the generalist pollinator *E. aeneus* solves this problem with a simple yet robust strategy of relying on a combination of visual and olfactory cues, rather than specializing on a single cue. We show that naïve hoverflies prefer multimodal visual and olfactory cues over unimodal cues (Fig. 2a.2, 2a.3).

Regarding visual cues, we found that colour choice in *E. aeneus* was mediated by brightness and not restricted to yellow hues. Floral objects reflecting in the 500-700 nm range (a yellow hue to human perception), were preferred over objects reflecting in the 400 – 500 nm range (Blue+o+), but in the absence of bright floral objects with yellow hues, hoverflies responded to Blue+o+ (Fig. 2b.4). This is similar to reports in other solitary pollinators like *Manduca sexta* (Kuenzinger et al. 2019). We also found that saturation (a limited range of wavelengths) was not a determining factor in response to floral objects. Naïve flies responded to floral objects reflecting across 400-700 nm (White+o+) as well as to objects with reflectance only in 500-700 (Fig. 2b.1). Thus, our data suggests that in flower naïve *E. aeneus*, response is not restricted specific hues or reflectance peaks when the floral objects contain odour. In addition, symmetry, but not disruptive shape, was also found important for object preference (Fig. 2c).

For olfactory cues, naïve hoverflies responded to several types and concentrations of odours indicative of plant life in general, such as green leaf volatiles and fruit volatiles, in addition to floral volatiles (Fig. 2d). However, we found that some odour cues were context-specific. Meta-cresol is a prominent odour of herbivore dung (Nair et al. 2018), a common oviposition site for female *E. aeneus*. Flower-naïve female flies preferred the positive control over a positive visual model paired with meta-cresol, but males responded to both models equally (Fig. 2d.4, 2d.5). Both male and female hoverflies have to forage for survival, hence foraging is not expected to be a sexually dimorphic trait in hoverflies. However, sex-specific foraging patterns have been observed in a few pollinators like the butterfly *Argynnis idalia idalia* (Balamurali et. al, 2020; Chmielewski et. al, 2023) and some bees (Roswell et. al, 2019) so we have analysed the floral object preferences of the hoverflies across sexes as well. Importantly, in our study meta-cresol was not repulsive to females, because naïve and mated female *E. aeneus* responded to meta-cresol in higher numbers when paired with a negative visual model (gray disk) than when paired with the positive visual model, although this result was not significant (Fig. 2e.1). Context matching and sexual dimorphism has been observed in other pollinators when foraging cues are juxtaposed with ovipositional cues (Balamurali et al. 2020). Thus, in specific contexts the same olfactory cue could trigger discrimination in females but not in males.

We conclude that the innate preferred nutritive object of *E. aeneus* consists of objects with radial symmetry, high brightness and plant odours (Fig. 2f, 4). It is probable that large regions of the peripheral olfactory and visual nervous systems detect these multiple cues rather than specialized neurons sensitive to specific floral features. As a consequence, innate flower identification in *E. aeneus* most likely results from an innate search template similar to global image matching found in bees, ants and wasps (Dittmar 2011), or snapshot matching in rodents for odour object identification (Gottfried 2010). Thus, the default state could be a lack of attraction until the right combination of cues exhibiting brightness, radial symmetry and plant odours are present. As a consequence, we hypothesize that the positive visual model with meta-cresol is not attractive to female *E. aeneus* (Fig. 2d) because the right combination of visual and olfactory cues is absent, rather than an aversion to the cues themselves.

To study how this proposed attraction to floral objects is processed in the hoverfly brain, response to these multimodal floral cues could be examined in the ventrolateral protocerebrum (vlPr). The ventrolateral protocerebrum (vlPr) is known to receive input from both the antennal lobe and the optic lobe in bees (medulla and lobula; Paulk et al. 2009; Liang et al. 2013) and inhibitory projection neurons necessary for attraction have been seen to suppress activity in vlPr in response to food odour (Strutz et al. 2014). The lateral horn (LH) of the fly brain has also been implicated in innate behaviour (De Belle and Heisenberg 1994). Interestingly, activity in the anterior-lateral LH, the region that is active in fruit flies exposed to odours of negative hedonic valence (Strutz et al. 2014), is mediated by odour-induced activity in the vlPr. Thus, the vlPR could be responsible for evaluating the right combination of visual and olfactory cues because when cues suppress activity in the vlPr, it has been shown to suppress aversive behaviour (Strutz et al. 2014).

By fractionating the object search template across multiple cues such as symmetry, brightness, and plant odour (Fig. 4), the hoverfly *E. aeneus* is capable of accepting objects innately with multiple perturbation in odours, colours, and shape, while still maintaining a robust search template. This simple multimodal gestalt can facilitate identification of multiple food objects in hoverflies, not only because these multimodal cues co-occur in specific objects in nature, but also because many of these cues indicate flower maturation and quality. In many flowering species, floral volatile emissions are maximized with flower maturation to increase pollinator visits (Schiestl et al. 1997; Muhlemann et al. 2006; Parachnowitsch et al. 2012). Pollination senescing flowers are fragrance-poor to avoid detection from seed predators (Dani et al. 2021). Conversely, intensification of petal colour is also observed in mature flowers with pollen (Schiestl et al. 1997). Further, previous studies show that disruption of radial symmetry by artificial manipulation and natural florivory deters pollinator activity (McCall 2008; Tsuji and Ohgushi 2018), with even small perforations reducing seed set (Tsuji and Ohgushi 2018). The innate preferences of *E. aeneus* for bright, radially symmetric objects with olfactory cues can function together as a gestalt that is flexible yet effective in identifying floral objects, allowing insects with tiny nervous systems to increase foraging efficiency without the need to increase neural processing.

**Fig. 4.**
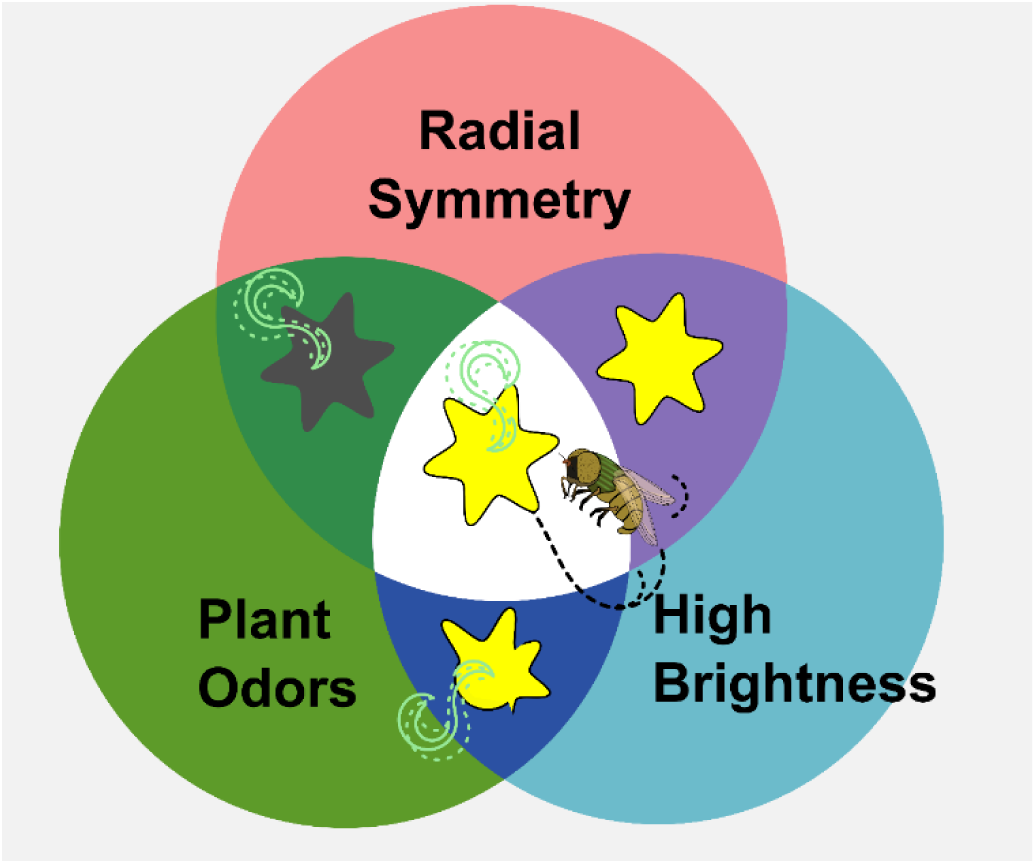
Schematic of the visual and olfactory cues important for preference to floral object in naïve *E. aeneus.* Objects with all three traits – high brightness, radial symmetry and plant odours are most attractive in our choice assays

## Acknowledgements

We would like to thank Roopa Rajendran, Deepa Rajan, Maansi Sharan for helping us with the maintenance and upkeep of the hoverflies. We are thankful to Subecha Rai and Dependra Rai for providing us with access to their farm for collection of the hoverflies and Dr. Malin Thyselius, and Dr. Olga Dyakova for adapting the rearing protocols for *E. aeneus*. We are also thankful to Dr. Gaiti Hassan, Dr. Axel Brockmann, Dr. Geetha G Thimmegowda, Hinal Kharva, Dr. Pavan K. Kaushik, Santosh Rajus, Dr. Cheyenne Tait, and Dr. Srishti Batra for their insights in different phases of the project and comments on the manuscript.

## Data accessibility

Supplementary data are provided in the electronic supplementary material.

## Authors’ contributions

A.M.: conceptualization, data curation, formal analysis, investigation, methodology, project administration, writing—original draft, writing—review and editing; A.J.: methodology, investigation; P.S.I.: investigation; A.S.: investigation; K.N.: conceptualization, funding acquisition, writing—review and editing. S.B.O.: conceptualization, funding acquisition, investigation, methodology, project administration, resources, software, supervision, validation, visualization, writing—original draft, writing—review and editing.

All authors gave final approval for publication and agreed to be held accountable for the work performed therein. No ethical approvals were required for our work and all required permissions were taken for the collection of specimens.

## Statements and Declarations

The authors have no competing interests to declare that are relevant to the content of this article.

**Supplementary_Information.zip contains the following**

**Online resource 1 contains the following:**

**Supplementary material S1:** Insect maintenance.

**Supplementary material S2:** Additional information on binomial GLMM used in the study.

**Fig. S1.** Illustration of experimental arena used in the study.

**Fig. S2.** Count versus gas chromatography retention index graphs for the odour blends used in the study.

**Fig. S3.** Dose-response curve of male versus female *E. aeneus* flies to meta cresol.

**Table S1.** Ratios of paints used in each floral model.

**Table S2.** Composition of volatiles used.

**Table S3.** Relative ratios of chemicals in the headspace of o+ blend compared to blend from Nordström et al, 2017, which was constituted from empirically measured release rates from flowers in nature as measured in the study.

**Table S4.** Number of flies tested in each assay, responding to either of the models in each assay (n_responders_), and responding to Model1 in each assay. P-value denotes whether the number of responders to Model1 is different from the number of responders to Model2. Number of flies tested for each assay and the response percentage

**Table S5.** Differences in response percentage between male and female flies across assays.

**Table S6.** Differences in preference index for model 1 between male and female responders across assays where a significant difference in response rates was observed between male and female flies.

**Table S7.** Differences in preference index for model 1 between male and female responders across assays without a significant difference in response rates between males and females.

**Table S8.** Differences in EAG response to doses of meta-Cresol treatment

**Table S9.** Tukey HSD post hoc test on the EAG responses to meta-cresol in male and female *E. aeneus*, n = 14 (males), n = 18 (females)

**Online resource 2**. Representative experiment trials depicting E. aeneus choosing between v+o+ and other morphs.

GCMS traces Folder, datafiles, and R scripts are available in Dryad (https://datadryad.org/stash/share/r1cnDo5enioFS5sGli5hbhWY6jv736BrbSC7Naxct9k**) as** follows –

**Dataset 1** – compilation of all normalised EAG responses recorded in the study.

**Dataset 2** – R scripts for GLMM, and analysis of sex difference in EAG responses to meta cresol.

**Dataset 3** – Compilation of all the observations undertaken in the study with information on response rate, sex, landings and visitations. This data is used in the GLMM described in supplementary methods S2.

**Dataset 4** – Reflectance spectra of models used in the study, and background of the experimental arena, graphed in Fig. 1.

