## SUPPLEMENTARY INFORMATION for "Innate floral object identification in the generalist solitary hoverfly pollinator *Eristalinus aeneus*"

**Online Resource 1: Supplementary Data File - This file contains Supplementary methods S1-S2, Fig. S1- S3, Tables S1-S9, and description for Online Resource 2, Dataset1, Dataset2, Dataset3 and Dataset 4.**

### SUPPLEMENTARY METHODS

#### S1. Insect maintenance

The *E. aeneus* used in the study were obtained with required permissions from three sources: Kalimpong, West Bengal, India from late May to early June 2018, Polyfly Almeria, Spain from August 2018 to January 2020, and the rest of the hoverflies were reared in the lab according to protocols established for *E. tenax* (Nicholas et al. 2018).

For lab rearing, third instar larvae were collected from manure pits and incubated to pupation in fresh cow dung in plastic boxes(Nicholas et al. 2018). Nearing pupation, the larvae were provided dry sawdust around the boxes and the larvae crawled into the sawdust for pupation. The sawdust was then sifted, and pupae were hand collected and transferred into petri plates with autoclaved vermiculite (Nicholas et al. 2018). The hatched adults were then transferred into aphid-proof nylon mesh cages of side 30 cm (width x height x length: 30 cm x 30 cm x 30 cm). The adult *E. aeneus* were supplied with distilled water, 10% sucrose solution, and ground mustard pollen mixed with powdered sugar *ad limitum* without exposure to flowers or plants. The volatiles in the headspace of ground mustard pollen were analysed using thermal desorption on PDMS (Polydimethylsiloxane), to ensure that none of the volatiles used in the current study could be detected in the pollen headspace chromatogram (Fig. S2). All *E. aeneus* were sex-sorted by day 4 unless the experiment required mated *E. aeneus*, and were housed in batches of 30 in chambers with temperature of 22$^{o}$C, humidity of 60%, and 12-hours light and 12-hours dark cycle. Only flower-naïve hoverflies between the ages of 3 - 14 days were used for experiments, and they were starved for a period of 4 - 6 hours prior to experimentation. The experiments were conducted in the day between 11 am and 6 pm.

#### S2. GLMMs used in the study.

For each assay, a series of GLMMs (Generalized Linear Mixed Models) were performed to test for differences in responses to the two models in each assay.

*Responder_Model1_Yes1_No0 ~ (1|Population),* [eq.1]

where, *Responder_Model1_Yes1_No0* denotes whether the fly responded to model1 in the assay or not (=1 if fly responded to model1; =0 if fly responded to model2), and

*Population* represented the batch from which the fly came, since multiple batches of flies were used in our experiments.

First, we tested whether the proportion of $n_{responder}$ flies responding to either of the models was different based on the sex of the fly (eq.2).

*Responder_Yes1_No0 ~ SEX_female1_male0 + (1|Population),* [eq.2]

where, *Responder_Yes1_No0* denotes whether the fly responded to either of the models in the assay, and

*SEX_female1_male0* denotes whether the fly is male or female (=0 if male; =1 if female).

If sex of the fly had an effect on whether it responded to either of the models (eq.2), we ran 2 separate GLMMs for each assay, one for only male flies and another only for female flies, testing for whether the $n_{responder}$ flies responded to the two models differently (eq.3a and eq.3b). We also separately tested for whether the proportion of $n_{responder}$ flies (irrespective of sex) were different in each assay (eq.3c).

*Responder_Model1_Yes1_No0 ~ (1|Population)*; only male $n_{responder}$ flies [eq.3a]

*Responder_Model1_Yes1_No0 ~ (1|Population)*; only female $n_{responder}$ flies [eq.3b]

*Responder_Model1_Yes1_No0 ~ (1|Population)*; all $n_{responder}$ flies [eq.3c]

If sex of the fly did not have an effect on whether it responded to either of the models (in eq.2), we tested for whether male and female $n_{responder}$ flies responded differently to the two models in the assay (eq.4a).

*Responder_Model1_Yes1_No0 ~ SEX_female1_male0 + (1|Population)*; [eq.4a]

In eq.4a, sex of the fly had an effect on response to model1 over model2 in only one assay (assay12; Supplementary Material Table S6). For the assays in which sex did not have an effect on one model over the other (eq.4a), we tested whether $n_{responder}$ flies (irrespective of sex) responded to the models differently (eq.4b):

*Responder_Model1_Yes1_No0 ~ (1|Population)*; all $n_{responder}$ flies [eq.4b]

For the assays in which very few males were involved (assay1, assay2, assay17, assay20), we tested whether the response was different to the two models among the $n_{responder}$ flies (eq.1). Note that we do not have data on the sex of the flies tested in model1=v+o+ vs. model2=v+1%GLV. Further, only gravid female flies were tested in the assay for model1=v+MC vs. model2=v-MC, and almost all female flies were tested in model1=v+o+ vs. model2=v-o- and model1=v+o+ vs. model2=v+o-

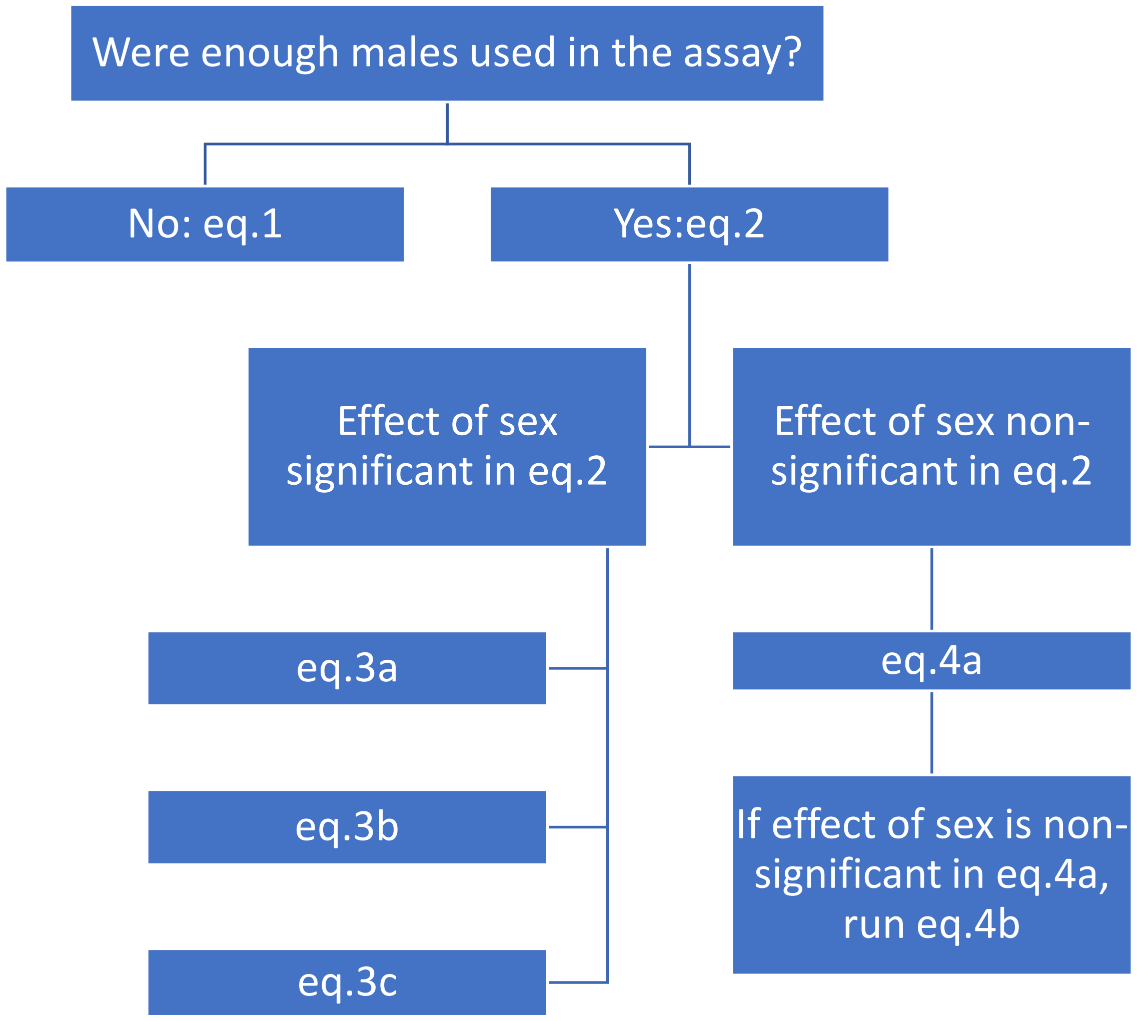
Were enough males used in the assay?
No: eq.1
Yes:eq.2
Effect of sex significant in eq.2
Effect of sex non-significant in eq.2
eq.3a
eq.3b
eq.3c
eq.4a
If effect of sex is non-significant in eq.4a, run eq.4b

### SUPPLEMENTARY FIGURES AND TABLES

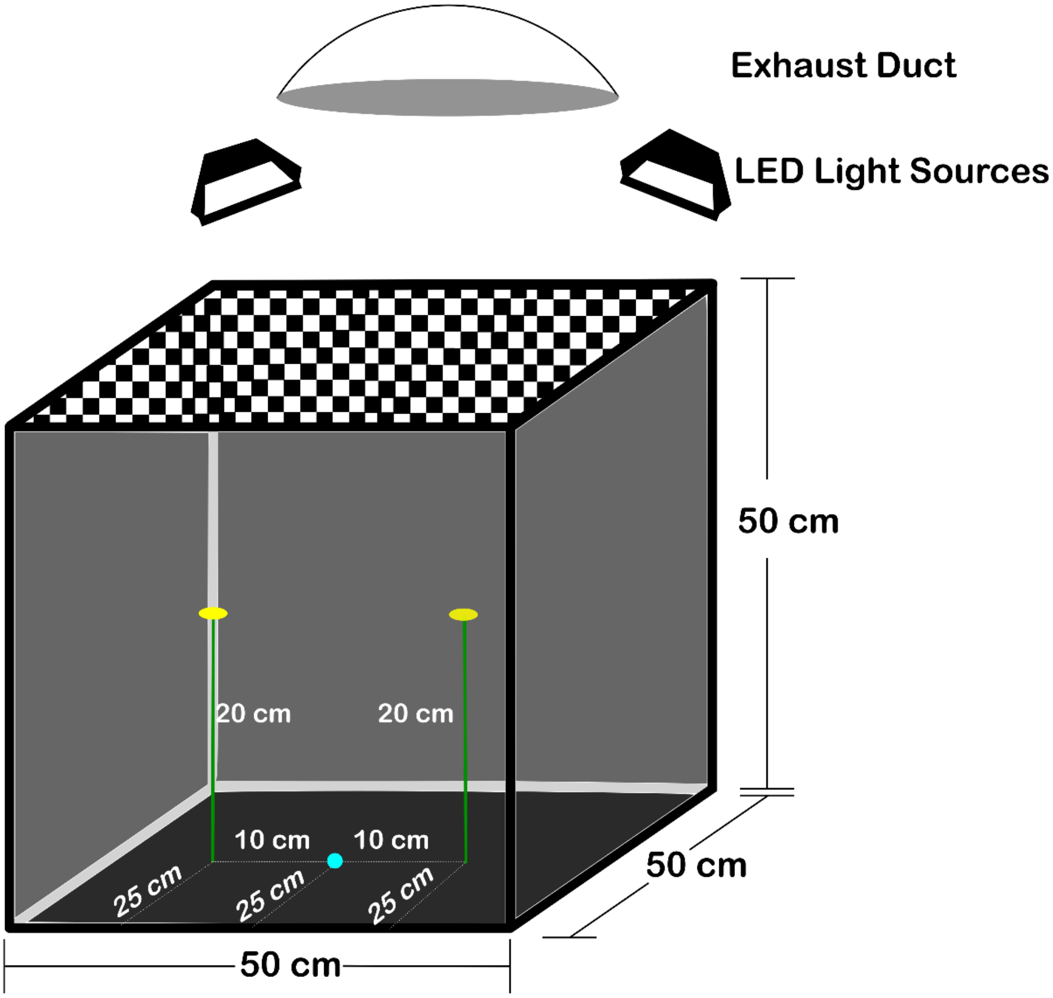

**Fig. S1** Illustration of the experimental arena used in the study. The blue circle on the base of the experimental arena indicates where *Eristalinus aeneus* hoverflies were released.

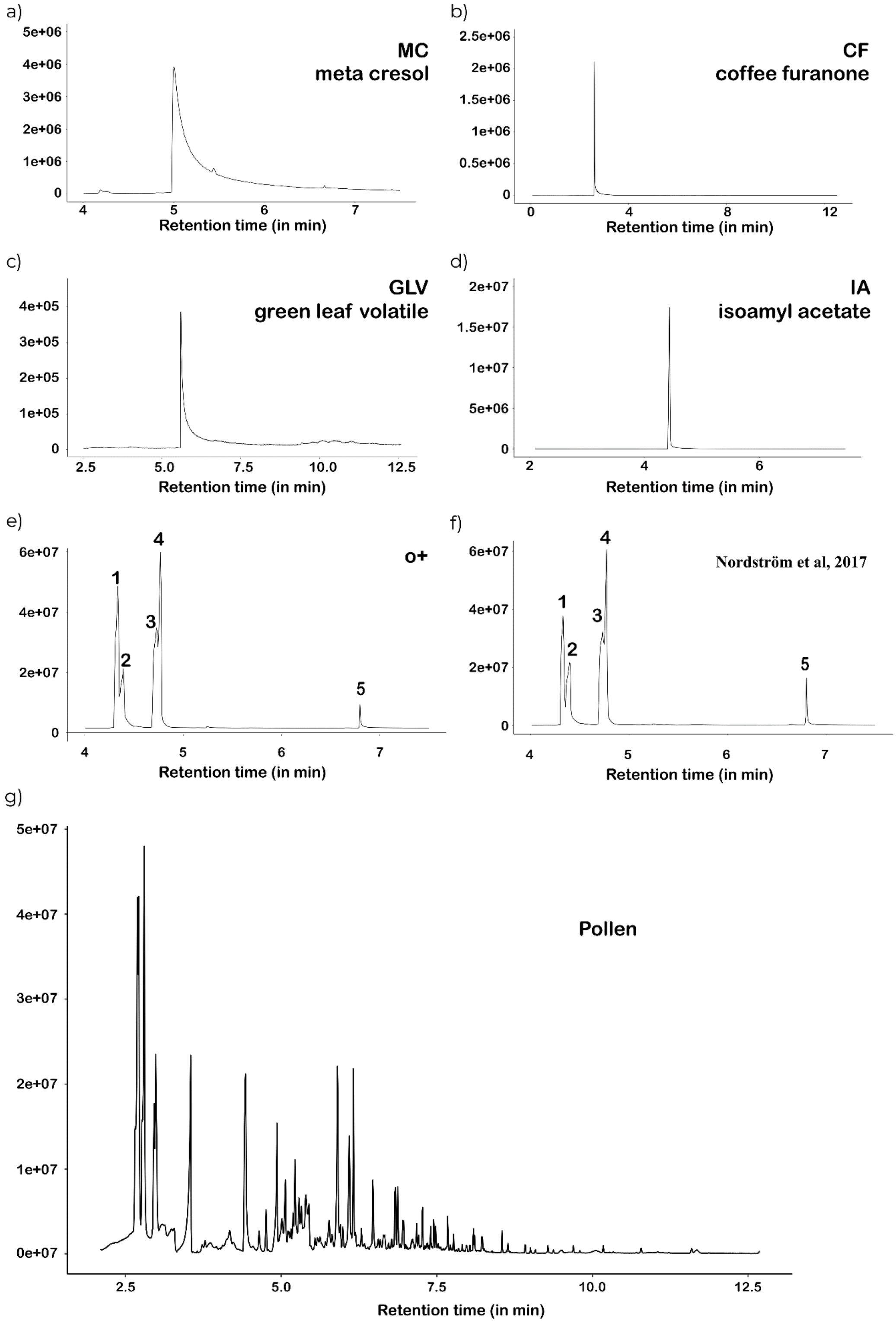

**Fig. S2** Gas chromatograms indicating retention time in minutes and abundance for the volatiles used in the study.

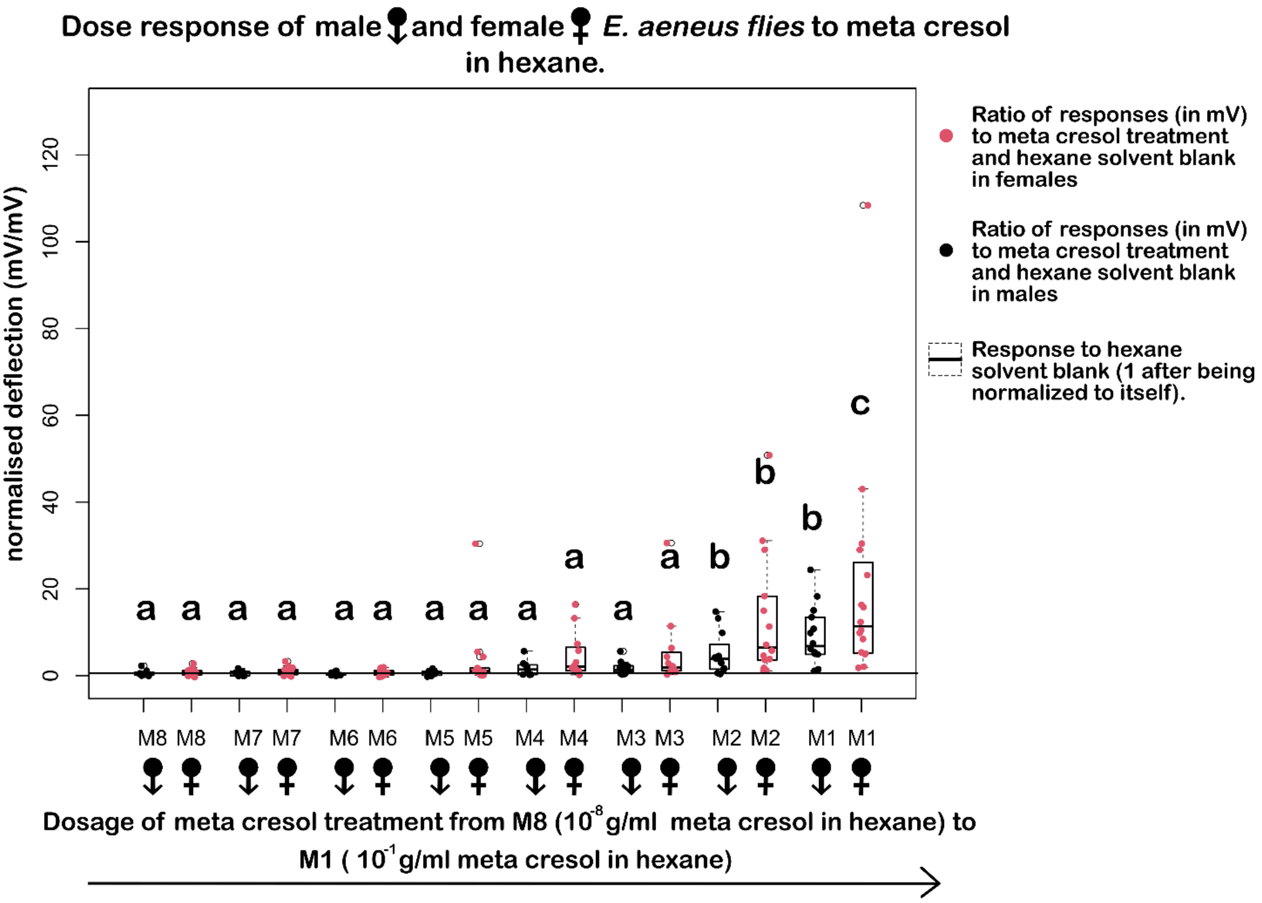

**Fig. S3** Dose-response curves of male and female *E. aeneus* flies to meta cresol.

**Table S1.** Ratios of paints used in each floral model.

| Floral model | Blend of Camlin acrylic colours |
|---|---|
| v+ /Asymmetric+/ Hexagon+ (all models have same paint as v+o+ i.e., yellow, reflectance peak 540-580 nm) | 14:1:3 ratio for Chrome yellow: sap green: white acrylic paint. |
| Low Reflectance LR+o+ (reflectance 540-580nm, reduced brightness compared to v+o+) | 14:1:3:1 ratio for chrome yellow: sap green: white: back acrylic paint. |
| Blue+ (reflectance peak 400-440 nm) | 100% Ultramarine blue acrylic paint. |
| White+ (reflectance peak 300-800 nm) | 100% White acrylic paint. |
| v- /Gray+ (reflectance peak 300-800 nm, reduced achromatic distance i.e., brightness compared to White+o+) | 9:1 ratio of White: black acrylic paint. |

**Table S2.** Composition of volatiles used.

| Odor blend | Composition |
|---|---|
| o+ (odour cues of v+o+) | 16.1 µl 2-Ethyltoluene [Sigma Aldrich, 99% purity]: 11.8 µl p-Cymene [Sigma Aldrich, 99% purity]: 60 µl Undecanal [Sigma Aldrich, 98% purity]: 24.1 µl r-Limonene [Sigma Aldrich, 97% purity]: 17.3 µl 6-Methyl-5-hepten-2-one [Fluka]: 82.7 µl mineral oil [Sigma Aldrich] |
| Coffee furanone (CF) | 10 µl 2-Methyltetrahydrofuran-3-one [Sigma Aldrich, 99% purity]: 90 µl mineral oil [Sigma Aldrich] |
| Vegetative odour – green leaf volatile (GLV) | 10 µl Cis-3-Hexenyl acetate [Sigma Aldrich, 98% purity]: 90 µl mineral oil [Sigma Aldrich] |
| Fruity odour-isoamyl acetate (IA) | 10 µl Isoamyl acetate [Sigma Aldrich, 98% purity]: 90 µl mineral oil [Sigma Aldrich] |
| Dung odor- meta cresol (MC) | 10 µl meta-Cresol [Sigma Aldrich]: 90 µl mineral oil [Sigma Aldrich] |
| 1%o+ | 100-fold dilution of o+ blend in mineral oil |
| 1%GLV | 100-fold dilution of GLV blend in mineral oil |

**Table S3.** Relative ratios of chemicals in the headspace of o+ blend compared to blend from Nordström et al, 2017, which was constituted from empirically measured release rates from flowers in nature as measured in the study

| **Blend** | **Retention Time** | **Area** | **Base peak** | **chemical** | **Relative ratio of areas under different chemicals of the blend** |
|---|---|---|---|---|---|
| **o+** | 4.33 | 101406628 | 105.1 | 2-Ethyltoluene | 27.59: 16.59: 53.16: 2.65 |
| | 4.39 | 60991256 | 108.1 | 6-Methyl-5-hepten-2-one | |
| | 4.77 | 195407267 | 93.1 | Limonene -Cymene | |
| | 6.8 | 9752319 | 57.1 | Undecanal | |
| **Blend from** | 4.32 | 77557758 | 105.1 | 2-Ethyltoluene | 22.21: 19.49: 53.48: 4.82 |

| | | | | |
|---|---|---|---|---|
| **Nordström et al. 2017** | 4.39 | 68037452 | 108.1 | 6-Methyl-5-hepten-2-one |
| | 4.76 | 186748875 | 93.1 | Limonene - Cymene |
| | 6.8 | 16831435 | 57.1 | Undecanal |

**Table S4.** Number of flies tested in each assay, responding to either of the models in each assay (**$n_{responders}$**), and responding to Model1 in each assay. P-value denotes whether the number of responders to Model1 is different from the number of responders to Model2. Number of flies tested for each assay and the response percentage

| **Assay** | **Number of flies tested** | **$n_{responders}$** | **Number of responders to Model1 (Preference index Model1)** | **Number of responders to Model2 (Preference index Model2)** | **P value** | **Probability of flies responding to Model1 [Mean (Lower 95% confint, Upper 95% confint)] irrespective of sex** | **Proportion of all flies tested responding to Model1** |
|---|---|---|---|---|---|---|---|
| **assay1; model1 =v+o+; model2 =v-o-** | 194 | 96 | 71 (73.96) | 25 (26.04) | $7.18*10^{-6}$ | 0.74 (0.64, 0.82) | 0.37 |
| **assay2; model1 =v+o+; model2 =v+o-** | 180 | 108 | 76 (70.37) | 32 (29.63) | $4.05*10^{-5}$ | 0.70 (0.61, 0.78) | 0.42 |
| **assay3; model1 =v+o+; model2** | 200 | 90 | 66 (73.33) | 24 (26.67) | $2.2*10^{-5}$ | 0.733 (0.631, .815) | 0.33 |

| | | | | | | | |
|---|---|---|---|---|---|---|---|
| **=v-o+** | | | | | | | |
| **assay4; model1 =v+o+; model2 = Gray+o+** | 82 | 56 | 41(73.21) | 15 (26.79) | 0.000862 | 0.732 (0.599, 0.833) | 0.5 |
| **assay5; model1= v+o+_L, model2= v+o+_R** | 190 | 103 | 50(48.53) | 53(51.46) | 0.768 | 0.485 (0.39, 0.582) | 0.26 |
| **assay6; model1 =v+o+; model2= Blue+o+** | 128 | 64 | 47 (73.44) | 17 (26.56) | 0.000327 | 0.734 (0.611, 0.83) | 0.37 |
| **assay7; model1 =v+o+; model2= v+GLV** | 118 | 61 | 29 (47.54) | 32 (52.46) | 0.701 | 0.475 (0.352, 0.602) | 0.25 |
| **assay8; model1 =v+o+; model2= White+o+** | 90 | 55 | 29 (52.73) | 26 (47.27) | 0.686 | 0.527 (0.394, 0.657) | 0.32 |
| **assay9; model1 =v+o+; model2= v+1%o+** | 110 | 62 | 40 (64.52) | 22 (35.48) | 0.0243 | 0.645 (0.517, 0.756) | 0.36 |
| **Assay10; model1 =v+o+; model2 =v+CF** | 120 | 60 | 44(73.33) | 16 (26.67) | 0.00053 | 0.733 (0.605, 0.831) | **0.37** |

| | | | | | | | |
|---|---|---|---|---|---|---|---|
| **assay11; model1 =v+o+; model2 =v+IA** | 108 | 58 | 28 (48.28) | 30 (51.72) | 0.696 | 0.475 (0.349, 0.604) | 0.26 |
| **assay12; model1 =v+o+; model2= *Asym+o+** | 100 | 61 | 46 (75.41) | 15 (24.59) | 0.00164 | 0.754 (0.628, 0.848) | 0.46 |
| **assay13; model1 =v+o+; model2 =LR+o+** | 54 | 30 | 24 (80) | 6 (20) | 0.00239 | 0.8 (0.611, 0.911) | 0.44 |
| **assay14; model1 =v+o+; model2= **Hex+o+** | 108 | 65 | 39 (60) | 26(40) | 0.181 | 0.582 (0.459, 0.696) | 0.36 |
| **assay15; model1 =LR+o+; model2= Blue+o+** | 100 | 64 | 32 (50) | 32 (50) | 1 | 0.5 (0.378, 0.622) | 0.32 |
| **assay16; model1= v+1%o+; model2= v+1%GLV** | 50 | 30 | 13 (43.33) | 17 (56.67) | 0.467 | 0.433 (0.264, 0.619) | 0.26 |
| **assay17; model1= v+MC; model2= v-MC** | 75 | 50 | 20 (40) | 30 (60) | 0.16 | 0.40 (0.27, 0.54) | 0.27 |

| | | | | | | | |
|---|---|---|---|---|---|---|---|
| **assay18; model1 =v+o+; model2 =v+MC (males)** | 192 | 126 | 65 (51.59) | 61 (48.41) | 0.469 | 0.602 (0.352, 0.808) | 0.34 |
| **assay18; model1 =v+o+; model2 =v+MC (females)** | 260 | 115 | 93 (80.87) | 22 (19.13) | $1.2*10^{-9}$ | 0.809 (0.725, 0.871) | 0.36 |
| **assay19; model1= v+1%o+; model2= v+GLV** | 113 | 61 | 25 (40.98) | 36 (59.02) | 0.161 | 0.41 (0.292, 0.539) | 0.22 |
| **assay20; model1 =v+o+; model2= v+1%GLV** | 84 | 57 | 34 (59.65) | 23 (40.35) | 0.148 | 0.596 (0.463, 0.717) | 0.4 |

* Asym+o+ is Asymmetric+o+, ** Hex+o+ is Hexagon+o+.

**Table S5.** Differences in response percentage between male and female flies across assays.

| **Assay** | **Intercept estimate; Logit scale (p-value for estimate in brackets)** | **Estimate for effect of sex; Logit scale (p-value for estimate in brackets)** | **Probability of flies responding of sex 0,1 0 = male, 1 = female, [Mean (Lower 95% confint, Upper 95% confint)]** | **Number of flies tested (number of responders) of sex 0,1 0 = male, 1 = female,** | **Number of flies tested (Number of responders) with sex unknown.** |
|---|---|---|---|---|---|
| assay3; model1=v+o+; | -0.1643 (0.523) | 0.3596 (0.319) | **0:** 0.459 (0.338, 0.585) | **0:** 61 (28) | 0 (0) |

| | | | | | |
|---|---|---|---|---|---|
| model2=v-o+ | | | **1:**<br>0.549 (0.456, 0.638) | **1:** 113 (62) | |
| assay4;<br><br>model1=v+o+;<br>model2=Gray+o+ | 0.3514 (0.2406) | 1.0349 (**0.0457**) | **0:**<br>0.587 (0.439, 0.721)<br>**1:**<br>0.8 (0.633, 0.903) | **0:** 46 (27)<br>**1:** 35 (28) | 1(1) |
| assay5; model1=v+o+_L, model2=v+o+_R | 0.0541 (0.849) | -0.05170 (0.848) | **0:** 0.513 (0.363, 0.649)<br>**1:** 0.501 (0.379, 0.622) | **0:** 64 (33)<br>**1:** 84 (42) | 43 (28) |
| assay6;<br><br>model1=v+o+;<br>model2=Blue+o+ | -0.3918 (0.349) | 1.4585 **(0.000401)** | **0:** 0.403 (0.228,0.608)<br>**1:** 0.744 (0.547, 0.875) | **0:** 73(26)<br>**1:** 52 (38) | 3 (0) |
| assay7;<br><br>model1=v+o+;<br>model2=v+GLV | -0.4568 (0.119269) | 1.8738 **(0.000136)** | **0:** 0.388 (0.261, 0.531)<br>**1:** 0.805 (0.653, 0.900) | **0:** 49 (19)<br>**1:** 41 (33) | 29 (9) |
| assay8;<br><br>model1=v+o+;<br>model2=White+o+ | 0.8329 (0.029) | -0.5101 (0.2828) | **0:** 0.697 (0.52, 0.83)<br>**1:** 0.580 (0.438, 0.71) | **0:** 33 (23)<br>**1:** 50 (29) | 7(3) |
| assay9;<br><br>model1=v+o+;<br>model2=v+1%o+ | -0.2272 (0.4696) | 1.0201 **(0.0198)** | **0:** 0.443 (0.299, 0.598)<br>**1:** 0.688 (0.539, 0.807) | **0:** 45(20)<br>**1:** 54 (37) | 12 (5) |
| assay10;<br><br>model1=v+o+;<br><br>model2=v+CF | 0.02985 (0.903) | -0.06759 (0.854) | **0:** 0.507 (0.388, 0.626)<br>**1:** 0.491 (0.358, 0.624) | **0:** 67 (34)<br>**1:** 53 (26) | 1(0) |

| | | | | | |
|---|---|---|---|---|---|
| assay11; model1=v+o+; model2=v+IA | -0.15414 (0.695) | 0.03637 (0.944) | **0:** 0.462 (0.281, 0.653)<br>**1:** 0.471 (0.309, 0.639) | **0:** 26 (12)<br>**1:** 34 (16) | 48 (31) |
| assay12; model1=v+o +; model2= Asymmetric+ o+ | 0.6495 (0.0397) | -0.4469 (0.2946) | **0:** 0.657 (0.506, 0.782)<br>**1:** 0.550 (0.381, 0.709) | **0:** 60 (39)<br>**1:** 40 (22) | 1(0) |
| assay13; model1=v+o +; model2=LR+ o+ | 0.4700 (0.244) | -0.4700 (0.395) | **0:** 0.615 (0.416, 0.782)<br>**1:** 0.500 (0.319, 0.681) | **0:** 26 (16)<br>**1:** 27 (14) | 0 (0) |
| assay14; model1=v+o +; model2= Hexagon+o+ | 0.43925 (0.220) | [-0.04513] (0.922) | **0:** 0.609 (0.432, 0.760)<br>**1:** 0.597 (0.419, 0.753) | **0:** 38(23)<br>**1:** 44(26) | 26(16) |
| assay15; model1=LR+ o+; model2= Blue+o+ | 0.8650 (0.0401) | [-0.2869] (0.5625) | **0:** 0.608 (0.432, 0.760)<br>**1:** 0.597 (0.419, 0.753) | **0: 27**(19)<br>**1:** 64(41) | 9(4) |
| assay16; model1= v+1%o+; model2= v+1%GLV | 0.51083 (0.323) | -0.08004 (0.898) | **0:** 0.625 (0.371, 0.825)<br>**1:** 0.606 (0.429, 0.759) | **0:** 33(20)<br>**1:** 16 (10) | 0(0) |
| assay18; model1=v+o +; model2=v+M C | 0.7932 (6.72^10- 07) | [-1.0250] **(4.24*10^ -07)** | **0:** 0.689 (0.618, 0.752)<br>**1:** 0.442 (0.383, 0.503) | **0:** 192 (126)<br>**1:** 260 (115) | 0(0) |
| assay19; model1= v+1%o+; model2= v+GLV | [-0.06643] (0.0788) | 0.72323 (0.125) | **0:** 0.483 (0.365, 0.604)<br>**1:** 0.659 (0.449, 0.820) | **0:** 83(40)<br>**1:** 26(17) | 4(4) |

144

**Table S6.** Differences in preference index for model 1 between male and female responders across assays where a significant difference in response rates was observed between male and female flies

| Assay | **Intercept estimate; Logit scale (p-value for estimate in brackets)**<br><br>**For males** | **Probability of flies responding to Model 1 [Mean (Lower 95% confint, Upper 95% confint)]**<br>**For males** | **Intercept estimate; Logit scale (p-value for estimate in brackets)**<br><br>**For females** | **Probability of flies responding to Model 1 [Mean (Lower 95% confint, Upper 95% confint)]**<br>**For females** |
|---|---|---|---|---|
| assay4; model1=v+o+; model2=Gray+o+ | 0.6931 (0.0895) | 0.667 (0.463, 0.823) | 1.5261 **(0.00198)** | 0.821 (0.625, 0.927) |
| assay6; model1=v+o+; model2=Blue+o+ | 0.47 (0.244) | 0.615 (0.41, 0.786) | 1.4881 **(0.000377)** | 0.816 (0.655, 0.912) |
| assay7; model1=v+o+; model2=v+GLV | 0.3080 (0.546) | 0.576 (0.317, 0.8) | 0.1823 (0.602) | 0.545 (0.37, 0.71) |
| assay9; model1=v+o+; model2=v+1%o+ | 0.4129 (0.397) | 0.602 (0.352, 0.808) | 0.7349 **(0.0366)** | 0.676 (0.505, 0.81) |
| assay18; model1=v+o+; model2=v+MC | 0.1313 (0.469) | 0.602 (0.352, 0.808) | 1.4416 **(1.2^10-09)** | 0.809 (0.725, 0.871) |

**Table S7.** Differences in preference index for model 1 between male and female responders across assays without a significant difference in response rates between males and females

| **Assay** | **Intercept estimate; Logit scale (p-value for estimate in brackets)** | **Estimate for effect of sex on preference index for model1; Logit scale (p-value for estimate in bracket)** | **Probability of male flies responding to Model 1 [Mean (Lower 95%** | **Probability of female flies responding to Model 1 [Mean (Lower 95%** |
|---|---|---|---|---|

| | | | confint, Upper 95% confint)] | confint, Upper 95% confint)] |
|---|---|---|---|---|
| Assay 3; model1=v+o+; model2=v-o+ | 0.7472 (0.0648) | 0.3949 (0.4312) | 0.679 (0.486, 0.825) | 0.758 (0.635, 0.85) |
| assay5; model1=v+o+_L, model2=v+o+_R | [-0.3054] (0.386) | 0.4964 (0.290) | 0.424 (0.267, 0.598) | 0.548 (0.395, 0.692) |
| assay8; model1=v+o+; model2=White+o+ | 0.4418 (0.301) | [-0.5108] (0.367) | 0.609 (0.397, 0.786) | 0.483 (0.307, 0.663) |
| assay10; model1=v+o+; model2=v+CF | [-0.33647] (0.566) | 0.08516 (0.912) | 0.417 (0.176, 0.705) | 0.438 (0.216, 0.687) |
| assay11; model1=v+o+; model2=v+IA | [-0.33647] (0.566) | 0.08516 (0.912) | 0.417 (0.176, 0.705) | 0.438 (0.216, 0.687) |
| assay12; model1=v+o+; model2=Asymmetric+o+ | 0.5798 (0.0824) | 2.4647 **(0.0221)** | 0.641 (0.478, 0.777) | 0.955 (0.730, 0.994) |
| assay13; model1=v+o+; model2=LR+o+ | 1.4663 (0.0221) | [-0.1671] (0.8549) | 0.812 (0.538, 0.942) | 0.786 (0.491, 0.933) |
| assay14; model1=v+o+; model2=Hexagon+o+ | 0.2624 (0.533) | 0.2076 (0.722) | 0.565 (0.358, 0.752) | 0.615 (0.415, 0.783) |
| assay15; model1=LR+o+; model2=Blue+o+ | [-0.1054] (0.819) | 0.1542 (0.781) | 0.474 (0.264, 0.693) | 0.512 (0.360, 0.662) |
| assay16; model1=v+1%o+; model2=v+1%GLV | [-1.3863] (0.0795) | 1.5870 (0.0810) | 0.20 (0.047, 0.559) | 0.55 (0.327, 0.755) |
| assay19; model1=v+1%o+; model2=v+GLV | [-0.51083] (0.118) | [-0.09531] (0.875) | 0.375 (0.238, 0.536) | 0.353 (0.165, 0.601) |

**Table S8.** Differences in EAG response to doses of meta-Cresol treatment

| **Doses being compared** | **diff** | **Lower bound** | **Upper bound** | **P adj** |
|---|---|---|---|---|
| p$10^{-7}$ and $10^{-8}$ | 0.17938028 | -10.6415126 | 11.00027 | 1 |
| $10^{-6}$ and $10^{-8}$ | 0.03273742 | -10.6726559 | 10.73813 | 1 |
| $10^{-5}$ and $10^{-8}$ | 2.17536790 | -8.6455249 | 12.99626 | 0.9986146 |
| $10^{-4}$ and $10^{-8}$ | 3.32933714 | -7.6166569 | 14.27533 | 0.9823644 |
| $10^{-3}$ and $10^{-8}$ | 3.91437675 | -6.7910166 | 14.61977 | 0.9511907 |
| $10^{-2}$ and $10^{-8}$ | 10.93617785 | 0.4371492 | 21.43521 | **0.0346344** |
| $10^{-1}$ and $10^{-8}$ | 17.48192212 | 7.3902143 | 27.57363 | **0.0000090** |
| $10^{-6}$ and $10^{-7}$ | -0.14664286 | -10.7240898 | 10.43080 | 1.0000000 |
| $10^{-5}$ and $10^{-7}$ | 1.99598762 | -8.6983407 | 12.69032 | 0.9991438 |
| $10^{-4}$ and $10^{-7}$ | 3.14995686 | -7.6709360 | 13.97085 | 0.9863363 |
| $10^{-3}$ and $10^{-7}$ | 3.73499647 | -6.8424505 | 14.31244 | 0.9594616 |
| $10^{-2}$ and $10^{-7}$ | 10.75679757 | 0.3882616 | 21.12533 | **0.0359678** |
| $10^{-1}$ and $10^{-7}$ | 17.30254184 | 7.3466640 | 27.25842 | **0.0000083** |
| $10^{-5}$ and $10^{-6}$ | 2.14263048 | -8.4348164 | 12.72008 | 0.9985452 |
| $10^{-4}$ and $10^{-6}$ | 3.29659972 | -7.4087936 | 14.00199 | 0.9810670 |
| $10^{-3}$ and $10^{-6}$ | 3.88163933 | -6.5776201 | 14.34090 | 0.9472334 |
| $10^{-2}$ and $10^{-6}$ | 10.90344043 | 0.6555009 | 21.15138 | **0.0282353** |
| $10^{-1}$ and $10^{-6}$ | 17.44918470 | 7.6189645 | 27.27940 | **0.0000048** |
| $10^{-4}$ and $10^{-5}$ | 1.15396924 | -9.6669236 | 11.97486 | 0.9999799 |
| $10^{-3}$ and $10^{-5}$ | 1.73900885 | -8.8384381 | 12.31646 | 0.9996279 |
| $10^{-2}$ and $10^{-5}$ | 8.76080995 | -1.6077261 | 19.12935 | 0.1657091 |
| $10^{-1}$ and $10^{-5}$ | 15.30655422 | 5.3506764 | 25.26243 | **0.0001316** |
| $10^{-3}$ and $10^{-4}$ | 0.58503961 | -10.1203537 | 11.29043 | 0.9999998 |
| $10^{-2}$ and $10^{-4}$ | 7.60684071 | -2.8921879 | 18.10587 | 0.3430947 |
| $10^{-1}$ and $10^{-4}$ | 14.15258498 | 4.0608771 | 24.24429 | **0.0007296** |
| $10^{-2}$ and $10^{-3}$ | 7.02180110 | -3.2261384 | 17.26974 | 0.4173878 |
| $10^{-1}$ and $10^{-3}$ | 13.56754537 | 3.7373252 | 23.39777 | **0.0009548** |
| $10^{-2}$ and $10^{-1}$ | 6.54574427 | -3.0593265 | 16.15082 | 0.4246962 |

**Table S9.** Tukey HSD post hoc test on the EAG responses to meta-cresol in male and female *E. aeneus*, n = 14 (males), n = 18 (females)

| p-value of the difference between | Male $10^{-2}$ g/ml m-Cresol | Male $10^{-1}$ g/ml m-Cresol |
|---|---|---|
| Female $10^{-2}$ g/ml m-Cresol | 0.7937453 | |
| Female $10^{-1}$ g/ml m-Cresol | | **0.0943278** |

**Online Resource 2.** Representative experiment trials depicting *E. aeneus* choosing between v+o+ and other morphs.

**GCMS traces Folder, datafiles, and R scripts are available in Dryad (https://datadryad.org/stash/share/r1cnDo5enioFS5sGli5hbhWY6jv736BrbSC7Naxct9k) as follows –**

**Dataset1 –** Compilation of all normalised EAG responses recorded in the study.

**Dataset 2 -** Rscripts for GLMM, and analysis of sex difference in EAG responses to meta cresol.

**Dataset 3 –** Compilation of all the observations undertaken in the study with information on response rate, sex, landings and visitations. This data is used in the GLMM described in supplementary material 1.

**Dataset 4 –** Reflectance spectra of models used in the study, background and lighting of the experimental arena, graphed in Fig. 1.

**References-**

Nicholas, S., Thyselius, M., Holden, M., & Nordström, K. (2018). Rearing and long-term maintenance of Eristalis tenax hoverflies for research studies. JoVE (Journal of Visualized Experiments), (135), e57711. 10.3791/57711
